# Drone Survey Reveals a Severe Chinstrap Penguin Decline and a Novel Gentoo Colony in an Antarctic Specially Protected Area

**DOI:** 10.1101/2025.10.15.682565

**Authors:** Eduardo J. Pizarro, Miguel Correa, Francine Timm, Gaspar Mejías, Ana Cláudia Franco, Juliana A. Vianna, Léa Cabrol, Francisco Santa Cruz, Lucas Krüger

## Abstract

Seabird populations in Antarctica are sensitive indicators of environmental change and understanding their dynamics is crucial for assessing the health and future of the Antarctic ecosystems. This study provides an updated population assessment for three key seabird species - the Antarctic Shag (*Leucocarbo bransfieldensis*), Gentoo Penguin (*Pygoscelis papua*), and Chinstrap Penguin (*P. antarcticus*) - within the Antarctic Specially Protected Area (ASPA) 133 on Nelson Island. Using drone-based imagery and photogrammetric analysis, we conducted comprehensive nest counts to evaluate current population size in comparison with historical data. Our results reveal a pronounced 57% decline in the Chinstrap Penguin population at Harmony Point since the 1990s, with 38,080 nests counted compared to historical estimates of nearly 90,000. In contrast, Gentoo Penguin numbers at Harmony Point (3,659 nests) appear stable following a previously documented increase. We also provide the first population estimate for a large Gentoo Penguin colony located on a small peninsula south of Harmony Point, known as the Toe, where 4,641 nests were recorded. Additionally, we report an 84% increase in Antarctic Shags, with 127 nests recorded at Harmony Point. These findings are consistent with regional species-level trends and provide critical baseline data for monitoring the impacts of rapid environmental change on Antarctic seabird communities.

## Introduction

The Maritime Antarctic Peninsula has experienced some of the most rapid warming in recent decades, leading to significant changes in sea ice dynamics, prey availability, and habitat conditions that affect seabird species (Morley et al. 2020; Ramos and Pereira 2022). Among these, the Antarctic shag (*Leucocarbo bransfieldensis*), chinstrap penguin (*Pygoscelis antarcticus*), and gentoo penguin (*Pygoscelis papua*) occupy important ecological niches, reflecting differences in foraging strategies, habitat use, and trophic dependence within the Antarctic marine ecosystem (Lynch et al. 2012; Lee et al. 2021). Gentoo penguins have demonstrated a notable capacity for range expansion and population growth, with increasing abundances and a southward shift in their distribution (Clucas et al. 2014; Herman et al. 2020). In contrast, chinstrap penguins have exhibited population declines across multiple regions, potentially linked to changes in krill availability and sea ice conditions (Trivelpiece et al. 2011; Strycker et al. 2020). In this context, Antarctic Specially Protected Areas (ASPAs) provide opportunities to evaluate long-term population trajectories under minimal direct human disturbance.

ASPA No. 133 on Nelson Island, South Shetland Islands, comprises two ice-free areas: Harmony Point and the Toe (Fig 1a-c). Harmony Point has long been recognized as an important breeding site for seabird species, especially chinstrap penguins (Silva et al. 1998). Although historical population data is available for some species, there are inconsistent gaps in monitoring data (Casaux and Barrera-Oro 2016; Oosthuizen et al. 2020), with the most recent penguin population estimates dating back to the mid-1990s (Silva et al. 1998). Against this background, although circumpolar satellite-based assessments have substantially advanced penguin monitoring, their ability to resolve some locations, including Nelson Island, has been limited by constraints in satellite imagery (Strycker et al. 2020). Drone technology and photogrammetric techniques therefore provide an effective complementary tool for more accurate and comprehensive seabird monitoring in remote and logistically inaccessible locations (i.e. Borowicz et al. 2018).

**Fig 1.**
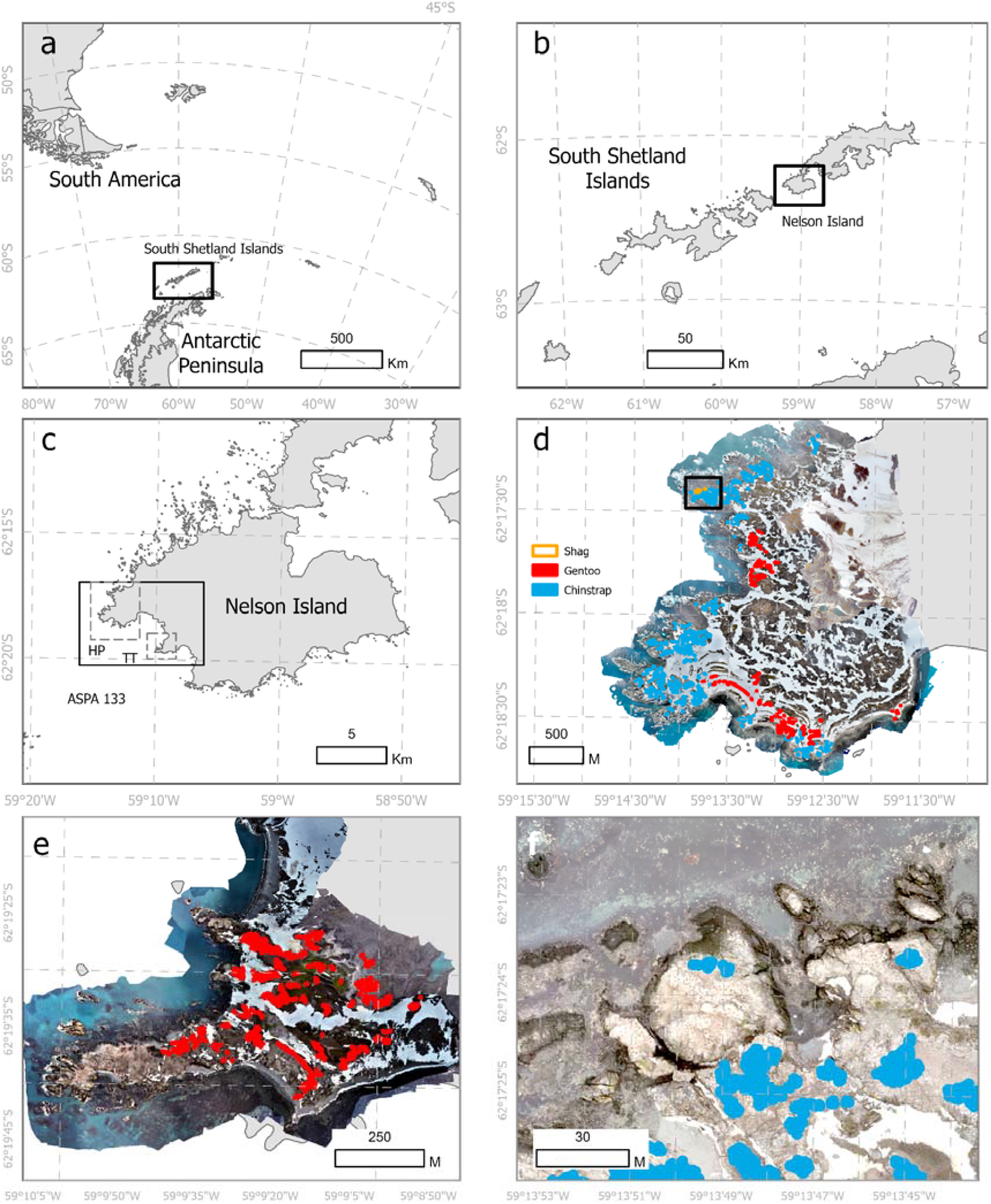
Location of the study area and distribution of seabird colonies. (a) Global context of the South Shetland Islands. (b) Position of Nelson Island within the Antarctic Peninsula. (c) Detail of Antarctic Specially Protected Area (ASPA) No. 133, showing the locations of Harmony Point (HP) and the Toe (TT) within the dashed squares. (d, e) Distribution of gentoo penguin (*Pygoscelis papua*) (red colour) and chinstrap penguin (*Pygoscelis antarcticus*) (blue colour) colonies at Harmony Point (d) and the Toe (e). (f) Distribution of Antarctic shag (*Leucocarbo bransfieldensis*) colonies at Harmony Point (orange colour). All distributions are mapped on high-resolution orthomosaics derived from drone imagery.

This study aims to provide an updated population assessment of Antarctic shags, chinstrap penguins, and gentoo penguins in ASPA No. 133, a protected area subject to minimal direct anthropogenic pressure, using drone-based imagery and photogrammetric analysis. By comparing current population estimates with historical data, we seek to contribute to the understanding of long-term population dynamics of these key indicator species within a rapidly changing Antarctic ecosystem.

## Methods

Aerial imagery was collected at the study sites Harmony Point and the Toe, using a DJI Mavic 3 unmanned aerial vehicle (UAV) equipped with a high-resolution camera (Fig 1a,b,c). Flights were conducted manually during favorable weather conditions at an elevation of 100 m above ground level. The theoretical ground sampling distance (following Stone and Davis 2025) was approximately 2.67 cm per pixel, according to camera specification: sensor width of 17.3, focal length of 12.29 and image width of 5280. Images were acquired at uniform resolution every 3s at an average speed of ∼5 m/s, resulting in a forward overlap of approximately 70–80% and side overlap of approximately 50-70%, while minimizing disturbance to breeding birds (Ratcliffe et al. 2015). At Harmony Point, a total of 7 flights were conducted over four survey days (6th, the 16th and 19th of December) during the incubation stage of gentoo and chinstrap penguins, yielding 5,932 images. These flights collectively covered the entire breeding area. At the Toe, two flights were conducted, one in 22th of December 2024, and another one on the 10th of January 2025 (to cover a small gap in the area detected after generating the orthomosaic) collecting 284 images. All drone imagery was processed using Agisoft Metashape Professional software v.2.2.1.20502 to generate one high-resolution orthomosaic image per flight. As orthomosaics were generated per flight, we calculated mean ± standard deviation resolution (distance between the center of consecutive pixels), x (longitude) and y (latitude) pixel size (in centimeters).

Flight operations adhered to the Environmental Guidelines for Operation of RPAS in Antarctica, adopted under Antarctic Treaty Consultative Meeting Resolution 4 (2018), and the most up-to-date handbook produced by COMNAP in collaboration with the SCAR committee COMNAP RPAS Operator’s Handbook (Version 8/18 December 2023), which provide best-practice recommendations for planning and conducting RPAS operations to minimize environmental impact in the Antarctic Treaty Area. Drone flights complied with all permit requirements for drone use in ASPAs (INACH permits Nº 00623/2024 and Nº 00624/2024).

Nest counts were conducted manually by a single observer placing a geographical point on each identified nest structure within the orthomosaic images using ArcGis Pro (ESRI 2026). The final shapefiles were used to estimate the number of nests. The identification of the Antarctic shags were corroborated by camera images taken on the field. The positions of gentoo penguin breeding groups at Harmony Point were marked on-site using a handheld GPS (Garmin Montana I7) to aid in species identification within the orthomosaic images. As we did not have access to the Toe site, the drone was launched from Harmony Point (∼ 3km). Although it is possible to distinguish between the two penguin species based on nest density, additional imagery was captured at an 80 m altitude to confirm species identification.

For Antarctic shags, a nest was defined as a seabird occupying a circular structure within the boundary of a white guano stain. For penguin species, a nest was considered occupied when an adult bird was observed either standing over or sitting on an identifiable nest structure within the boundary of a guano stain, and forming part of a recognizable breeding aggregation (i.e., multiple nests in close proximity, consistent with colony structure). For estimating accuracy of the counts, three different authors (EP, GM and LK) marked nest points on a subset of spatially discrete units of the colonies for chinstrap (n=38) and gentoo (n=24), and all of the colony for the shags (n=10). The difference between the three counts was used to calculate standard deviation, and then the accuracy calculated as the percentage of the standard deviation corresponding to the mean of the three counts (coefficient of variation). We also calculated the coefficient of variation to the total of the counts per observer, in order to compare whether differences in counts at smaller subsets reflects the differences in total count. We tested whether there was correlation between counts using a two-way test intraclass correlation coefficient in the ‘irr’ R package (Matthias Gamer et al. 2025). The topography of Harmony Point was estimated from previous drone flights (Santa Cruz and Krüger 2023).

Current population estimates were compared with historical data from previous studies conducted at Harmony Point. For Antarctic shags, population estimates are available from (Casaux and Barrera-Oro 2016) and (Oosthuizen et al. 2020), while comparable information for chinstrap and gentoo penguins is provided by (Silva et al. 1998) and (Croxall and Kirkwood 1979). Early population assessments were largely based on coarse visual surveys and extrapolations and are generally considered to have high uncertainty, often on the order of several tens of percent (e.g., Croxall and Kirkwood 1979). In contrast, drone-based surveys, such as those used in the present study, offer substantially higher spatial resolution and improved methodological consistency. Although the accuracy of drone-derived estimates varies depending on image resolution, survey design, and counting protocols, they are generally expected to provide lower uncertainty than early ground-based assessments (Humphries et al. 2017; Hilton et al. 2024).

All analyses were conducted in R (R Core Team 2024) and maps generated in ArcGIS Pro (ESRI 2026). All data and R code is available at Krüger et al. (2026) and (Krüger 2026). Inter-Quartil Range (IQR) are presented as lower and upper 75%.

## Results

The resulting orthomosaics had 10.25 ± 1.16 cm/pix of resolution (min 9 cm, max 12 cm). X pixel size was 7.30 ± 0.83 cm (min 6.20 cm, max 8.62 cm). Y pixel size was 3.40 ± 0.39 cm (min 2.89 cm, max 4.01 cm).

Median accuracy of the counts was 5.1% [IQR 2.3%, 8.3%], 4.7% [2.6%, 7.5%] and 4.5% [0.9%, 6.5%] for chinstrap penguins, gentoo penguins and Antarctic shags, respectively. Total variation was within the IQR for chinstrap penguins (4.3%) and Antarctic shags (1.2%) but was below the IQR for gentoo penguins (1.9%). Very high agreement was found between observers for the three species: chinstrap penguin ICC = 0.994 [IQR 0.988,0.997], F_1,37_=698.4, p<0.001; gentoo penguin ICC = 0.993 [0.988,0.997], F_1,19_=511.8, p<0.001; Antarctic shag ICC = 0.949 [0.863,0.986], F_1,9_=53.7, p<0.001.

Chinstrap penguins occupied most of the areas along the shoreline and rocky outcrops (Fig 1d,e). A total of 38,080 and 8,153 nests were recorded in the 2024 season at Harmony Point and the Toe, respectively. This represents a decrease of 57% and 74% from the 1990s and 1980s estimates respectively, for Harmony Point (Fig 2a). For the Toe colony, it represents a 45% decrease from the 1970s, with a high error around that estimation (±50%) (Fig 2b). Chinstrap penguins occupied elevations mostly between 9.8 m and 32.9 m, on slopes of 1.8º to 5.6º inclination, and were typically found within 104 m of the coastline.

**Figure 2.**
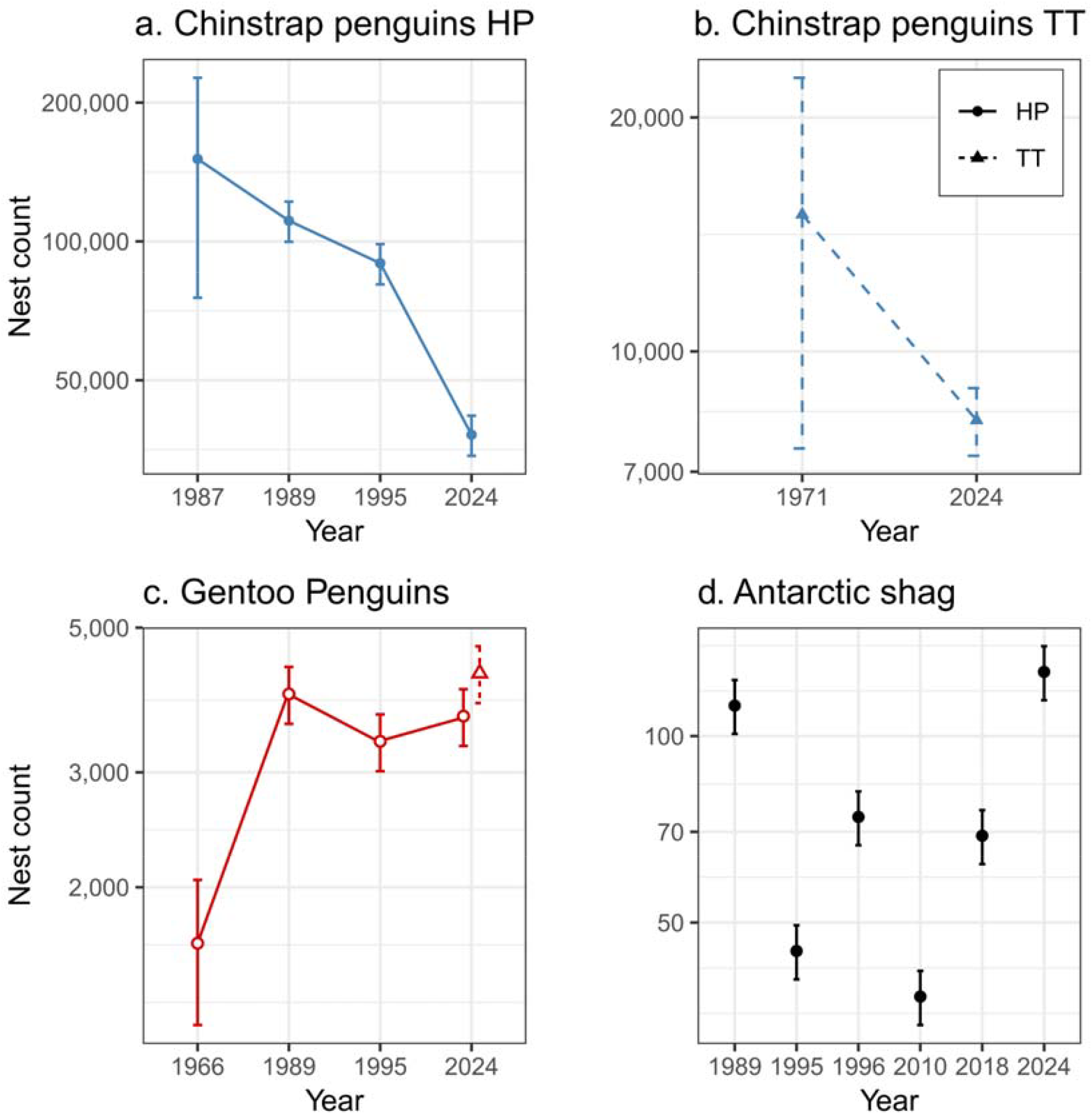
Colony size ± 10% or ± 50% (for penguins counts before 1985) error for chinstrap penguin (a,b), *Pygoscelis antarcticus* (blue filled shapes), gentoo penguin (c), *Pygoscelis papua* (red empty shapes), and Antarctic shag (*Leucocarbo bransfieldensis*) (d), measured as number of nests, in Harmony Point (HP, circle) and the Toe (TT, triangles), ASPA 133, Nelson Island. Year 2024 corresponds to data from this study, as previous years are from previous publications (please see methods section).

Gentoo penguins nests were estimated at 3,659 and 4,641 in the 2024 austral summer season at Harmony Point and the Toe, respectively (Fig 1d,e). Previous assessments indicate that the Harmony Point colony has fluctuated between 3,000 and 4,000 pairs since the late 1980s, with one earlier estimate of 1,642 pairs in 1966 (Fig 2c), representing an 122% increase since the 1960s. Most nests were located at elevations between 10.3 m and 36.0 m, on smooth slopes with inclinations of 1.0º to 5.4º, and at distances of 67.3 m to 151.7 m inland.

A total of 127 Antarctic shag nests were recorded at Harmony Point (Table 1, Fig 1f). This represents an 84% increase compared to the previous assessment in the 2018/19 breeding season but falls within the expected variability considering counts between 1980 and 2010 (Fig 2b). Compared to the penguin colonies, shags occupied lower elevations (11.2 m to 17.0 m), on relatively steeper slopes (10.4º to 18.4º), and closer to the coastline (within 15 m inland).

## Discussion

We update the population size and spatial distribution of *Pygoscelis* penguins in ASPA No. 133, which had not been revised since the 1990s. The population assessment demonstrates a marked decline of 57% in one of the largest chinstrap penguin colonies in the South Shetland Islands, consistent with recent regional assessments (Strycker et al. 2020; Hilton et al. 2024; Hinke et al. 2025), and report the presence of a previously undescribed, large gentoo penguin colony at the Toe with over 4,500 pairs, contributing to further evidence of population expansion in this species (Herman et al. 2020). Together, these findings highlight the importance of site-based population assessments and the value of drone-derived data for accurately quantifying breeding populations in remote Antarctic environments.

The number of chinstrap penguins at Harmony Point exhibited a clear decline from the last available estimate of nearly 90,000 breeding pairs in the mid-1990s (Silva et al. 1998). The observed 57% decrease over three decades is consistent with regional declines reported across the Antarctic Peninsula (Talis et al. 2023; Krüger 2023; Hinke et al. 2025), and with assessments suggesting increasing conservation concern for the species at a local scale (Krüger 2023). Many chinstrap colonies in the region still lack recent and reliable population estimates (Harris et al. 2015), highlighting the importance of updated, site-specific assessments such as those presented here. The concordance between the decline observed at Harmony Point and broader regional trends supports the robustness of this result. Although the underlying causes remain poorly understood, several interacting factors are potentially affecting chinstrap penguin population dynamics, include the combined effects of climate change on chick survival (Salmerón et al. 2023), shifts in prey availability combined with increasing krill fisheries (Watters et al. 2020; Krüger et al. 2021), and competition with recovering whale populations (Savoca et al. 2024). We hypothesize that the cumulative effects of multiple factors are likely responsible for the observed trends. The Maritime Antarctic Peninsula holds one-third of the global chinstrap penguin population (Strycker et al. 2020); even assuming the remaining two-thirds of the population is stable, the observed decline suggests a substantial global reduction.

In contrast, gentoo penguin numbers at Harmony Point increased between the 1960s and the 1990s and appear to have stabilized in recent decades (Croxall and Kirkwood 1979; Silva et al. 1998). As a species with greater dietary flexibility than other pygoscelids, gentoo penguins have demonstrated a higher capacity to cope with environmental change in the region (Miller et al. 2009; McMahon et al. 2015), allowing them to maintain or expand populations while sympatric species decline (Talis et al. 2023). For the Toe colony, where no previous data was available, our results serve as a baseline for future trend estimations.

Historical data for Antarctic shags from 1964 to the present indicate substantial fluctuations in colony size over time, with the current estimate of 127 nests representing the highest value recorded to date, pattern that is consistent with previous studies reporting high spatial and temporal variability in Antarctic shag populations throughout the Antarctic Peninsula region (Casaux and Barrera-Oro 2006, 2016; Schrimpf et al. 2018).

Our results are consistent with previously reported nesting habitat segregation among penguin species, primarily structured by distance inland and elevation (Gallagher et al. 2025; Korczak□Abshire et al. 2026), with gentoo penguins occupying more inland nesting areas than chinstrap penguins. This reflects species-specific nesting preferences and the greater capacity of gentoo penguins to exploit a broader range of ice-free habitats, as previously reported along the West Antarctic Peninsula (Cole et al. 2019; Herman et al. 2020). The preferential use of inland areas by gentoo penguins may facilitate colonization of newly exposed habitats as ice-free land expands under ongoing regional warming (Lee et al. 2022). Such habitat segregation has important implications for future species distributions and potential interspecific interactions in rapidly changing Antarctic coastal ecosystems.

Satellite imagery provides a valuable broad-scale framework for monitoring seabird populations; however, its application at local scales is constrained by image resolution, atmospheric conditions, and classification uncertainty (Román et al. 2022; Hilton et al. 2024). Using medium-resolution satellite imagery from 2014, (Strycker et al. 2020) estimated 3,000 and 10,000 nests at Harmony Point and the Toe, respectively, whereas our drone-based survey recorded substantially higher nest numbers at both sites. The differences found are likely due to methodological limitations of satellite data rather than true population change, since Strycker et al. (2020) could not retrieve good quality images for those sites. Together, satellite imagery and UAV surveys provide complementary information especially in remote areas, with satellite data supporting broad-scale monitoring and drone-based assessments delivering the fine-scale accuracy required for reliable population models and future seabird assessments.

ASPAs such as Harmony Point play a crucial role in conserving seabird hotspots by providing protected refuges where direct human disturbances, including tourism and research station infrastructure, are minimized. These areas contribute to local ecosystem stability and the maintenance of species interactions within sensitive Antarctic habitats, as shown for benthic communities (Moya et al. 2025). However, ASPAs alone cannot mitigate large-scale threats such as the ongoing warming, pollution, climate change and fisheries, which affect seabirds at regional scales. Therefore, maintenance of seabird populations requires conservation strategies that complement local protection with broader, ecosystem-based management approaches to safeguard Antarctic biodiversity.

## Statements and Declarations

### Data Availability Statement

Data is available in repository (Krüger et al. 2026).

## Acknowledgments

We thank the crew of the vessel Karpuj for transportation to and from the study site, and INACH logistics for their support in organizing the expedition.

